# Knockdown of DJ-1 Resulted in a Coordinated Activation of the Innate Immune Antiviral Response in HEK293 Cell Line

**DOI:** 10.1101/2024.06.20.599923

**Authors:** Keren Zohar, Michal Linial

## Abstract

PARK7, also known as DJ-1, plays a critical role in protecting cells by functioning as a sensitive oxidation sensor and modulator of antioxidants. DJ-1 acts to maintain mitochondrial function and regulate transcription in response to different stressors. In this study, we show that cell lines vary by their antioxidation potential at basal condition. The transcriptome of HEK293 cells was tested following knockdown (KD) of DJ-1 using siRNAs which reduced the DJ-1 transcripts to only 12% of the original level. We compared the expression levels of 14k protein coding transcripts, and 4.2k non-coding RNAs relative to control cells treated with non-specific siRNAs. Among the coding genes, ∼200 upregulated differentially expressed genes (DEGs) signified a coordinated antiviral innate immune response. Most genes were associated with regulation of type 1 interferons (IFN) and induction of inflammatory cytokines. About a quarter of these genes were also induced in cells treated by non-specific siRNAs that were used as a negative control. Beyond the antiviral response, 114 genes were specific to KD of DJ-1 with enrichment in RNA metabolism and mitochondrial functions. A smaller set of downregulated genes (58 genes) were associated with dysregulation in membrane structure, cell viability, and mitophagy. We propose that KD of DJ-1 diminish its protective potency against oxidative stress, rendering the cells labile and responsive to dsRNA signal by activation of a large number of genes, many of which drive apoptosis, cell death, and inflammatory signatures. The KD of DJ-1 highlights its crucial role in regulating genes associated with antiviral responses, RNA metabolism, and mitochondrial functions, apparently through alteration in STAT activity and downstream signaling. Given that DJ-1 is highly expressed in metastatic cancers, targeting DJ-1 could be a promising therapeutic strategy where manipulation of DJ-1 level may reduce cancer cell viability and enhance the efficacy of cancer treatments.

## Introduction

Parkinson’s disease (PD) is a neurodegenerative disorder characterized by the progressive loss of dopaminergic neurons, resulting in motor dysfunction and cognitive impairment. Neuronal cell death under oxidative stress and mitochondrial dysfunction are considered major drivers of PD pathologies [1]. DJ-1’s primary role as an essential mitochondrial protein explains much of its role in PD [2]. Mutations in PARK7 (also called DJ-1) were implicated in familial early-onset and sporadic forms of PD [3]. As an oxidative sensor, DJ-1 plays a critical role in maintaining redox homeostasis in all cells by sensing and modulating cellular oxidative responses. DJ-1 is involved in oxidation stress in direct and indirect modes. For example, DJ-1 binds the subunits of mitochondrial complex I. Therefore, upon DJ-1 depletion, mitochondrial function is drastically reduced. Downregulating DJ-1 in neuronal cells led to increased cell death in response to oxidative stress, ER stress, and proteasome inhibition, while overexpression of DJ-1 has rescued cell death [4]. Several oxygen-based modifications to the cysteine residues of DJ-1 create a set of modified variants that are partitioned among cellular compartments (mitochondria, cytosol, nucleus, and exosomes) to conduct downstream functions in signaling and transcription regulation [5]. To protect cell death from various stressors, the modified protein is engaged in a rich protein interaction network that includes often specific signaling pathways and cell types [6, 7].

The DJ-1 has garnered considerable interest due to its multifaceted roles in driving numerous processes [1, 8]. DJ-1 is involved in transcriptional regulation in direct and indirect modes. To gain insights on the role of DJ-1 in cells, diverse cell systems were used. This includes PD-related cell models from humans and rodents, such as primary dopaminergic neurons [9], brain slices, and numerous cell lines [10]. Expanding the role of DJ-1 in cell signaling, argue that its location, and the pattern of its post-translational modifications determine the effect of DJ-1 in different cell systems [7], a studied by overexpression [11] or knockdown (KD) (e.g., [12, 13]).

Accumulating evidence validates the role of DJ-1 as an oncogene in cancer biology and, more specifically, in tumorigenesis and cancer progression [14]. The expected impact of DJ-1 on cancer is tightly connected to its role in apoptosis, cell proliferation, and metastasis. For example, the role of DJ-1 in suppressing apoptosis in cancer cells allows for the production of the driving oncoproteins, thereby promoting tumor progression. However, the molecular mechanisms underlying DJ-1-mediated modulation of cancer-related pathways remain elusive. In cancer cells, DJ-1 regulates the JNK pathway and consequently impacts autophagy [15, 16]. This was further confirmed in prostate cancer [17]. DJ-1, through the regulation of androgen receptor (AR) signaling, affected Beclin1-involved autophagy responses via the JNK-dependent pathway. DJ-1 overexpression reduces LC3 transformation and autophagosome formation, while DJ-1 knockdown has the opposite effect. JNK phosphorylation and Bcl2 dissociation are affected by DJ-1 levels. Recent studies have highlighted the transforming activity of cancer cells. For example, together with H-Ras/Myc, the protein translocates from the cytoplasm to the nucleus during the S phase of the cell cycle. Overexpression of DJ-1 enhances colorectal cancer cell proliferation through the cyclin-D1/MDM2-p53 signaling pathway [18] and acts to increase cancer cell survival through the PI3K-AKT pathway [19]. The elevated levels of DJ-1 have been detected in breast, lung, and prostate cancers. making it a potential biomarker for diagnosis and prognosis [14, 20].

DJ-1’s function in the brain has shown its involvement in controlling inflammation [21]. Higher levels of inflammation signatures (including TNF, iNOS, NO, IL-6, cyclooxygenase-2, and p38 phosphorylation) were observed in astrocytes and microglia lacking the DJ-1 mouse model after lipopolysaccharide (LPS) exposure [22]. Similarly, brain slices from DJ-1 knockout mice exposed to IFN-γ exhibited increased levels of IL-6 and TNF, as well as STAT1 phosphorylation [23]. Studies that mimicked glial activation by injecting LPS into the substantia nigra of DJ-1 knockout mice resulted in elevated levels of immune-responsive genes including ICAM-1, IFN-γ, IL-1β, IL-16, IL-17, CXCL11, and more [24]. Reducing DJ-1 levels also accelerated IKK and IkBα phosphorylation, leading to enhanced p65 nuclear translocation in both normal and LPS-exposed cells [6]. The presence of elevated NF-κB promoter function in DJ-1-depleted cells further confirms DJ-1’s inhibitory function in neuroinflammation [25].

Upon viral infection, the innate system is activated leading antiviral defense mechanism mediated by interferon (IFNs). The type I IFNs (IFNα, IFNβ) activate IFN receptors (IFNARs) via JAK1, TYK2, STAT1, and STAT2 signaling pathways, whereas type II IFN (IFNγ), activates IFNγ receptors (IFNGRs), leading to STAT1 phosphorylation and gene expression. Upon viral infection (e.g., Respiratory syncytial virus; RSV), reactive oxygen species (ROS) levels increase and as a result of changes in cell redox the JAK-STAT pathway is activated, resulting in expression of antiviral genes [26, 27]. Elevated ROS signifies the pathogenesis of human diseases such as pulmonary fibrosis, Parkinson’s and Alzheimer’s diseases [28].

In this study, we tested the impact of changing the naïve level of DJ-1 by suppress its expression in HEK293 cells. We performed RNA-seq global analysis to assess the role of knockdown using siRNA for DJ-1 levels. We show that siRNA causes a stress response that was converted to antiviral cascade. The lack of DJ-1 in cells causes a failure in redox homeostasis and coordinated activation of the STAT pathways that connect a transcriptional wave of interferon-based antiviral response and a direct effect on mitochondria and RNA metabolism. The potential role of targeting DJ-1 in activating the antiviral response in metastatic cancers is discussed.

## Methods

### 2.1. Cell lines and siRNAs

Human neuroblastoma SH-SY5Y cells (CRL-2266) were purchased from ATCC (American Type Culture Collection, Rockville, MD, USA). Cells were cultured in Minimum Essential Media (MEM and F12 ratio 1:1, 4.5 g/L glucose) with 10% fetal calf serum (FCS) and 1:10 L-Alanyl-L-Glutamine. Cell cultures were incubated at 37 °C in a humidified atmosphere of 5% CO2.

HEK293 cells were cultured to reach a 20-30% confluent level (3× 10^4^ per cm2). For transfection, we have used Lipofectamine (Gibco-Invitrogen) in cells at 40-50% confluency according to the manufacturer’s instructions. Lipofectamine 2000 was shown to be ideal for HEK293 [29]. The esiRNA system (Invitrogen Catalog #: 11668019) consists of a pool of hundreds of siRNA (21 bp each), where each individual dsRNA has a low concentration in the pool which diminish most off-target effects, while producing an efficient knockdown. We apply the esiRNA-RULC of a mixture of 21 nt dsRNA used as a negative control (RLUC stands esiRNA directed against Renilla Luciferase; EHURLUC) in addition to the specific PARK7 (to knockdown the DJ-1 transcripts and protein products). RULC experiments were used to measure the baseline effects of introducing esiRNA that controls for the delivery method and allows to distinguish the cellular response to the targeted siRNA itself [30].

### 2.2. Cell staining by immunofluorescence

The SH-SY5Y cells were seeded and grown in 24-wells plate containing slide discs coated with Collagen solution (2% Acetic, 2% Collagen, 96% DDW). After 24 h, cells were washed (x2) in PBS at room temperature (RT) and cells were fixed by freshly prepared 4% paraformaldehyde (PFA in PBS, at RT). Following 10 min incubation, the slides were then washed (x3) and incubate for 15 min with 0.1% Triton X-100 in PBS. The perforated cells were incubated for 60 min at RT in blocking solution with 1% BSA in PBS, drained and incubated with DJ-1 primary antibody diluted 1:500 in 1% BSA in dark -humid chamber for 3 h (or overnight, 4 °C) followed by washing and staining by secondary antibody (1:5000) for 45 min at RT (keep cells dark in a humidified chamber). We stained the nuclear with DAPI (1μl/ml, 5 min). Slides were quenched using sodium borohydride (NaBH4, 1 mg/ml) in PBS for 5 min and mounted on slides for microscopy. All procedures with fluorescence antibodies and mounting protocol were performed in a dark-humid chamber. Washing was performed with PBST with 3 consecutive times, 5 min each between steps.

### 2.3. Reverse transcription PCR (RT-PCR) & PCR

Cells were collected for RNA preparation. Total RNA was extracted from cell culture with TRIzol (Thermo-Fisher Scientific, Waltham, MA, USA), and RT was performed using a Ready-To-Go first-strand synthesis kit (Cytiva, Marlborough, MA, USA) according to the manufacturer’s instructions. RNA was reverse transcribed into cDNA (1 μg) and used in the PCR reaction. The PCR conditions consisted of denaturation at 95 °C for 2 min and 35 cycles (10 s at 95 °C, 15 s at 60 °C, 30 s at 72 °C), and 5 min for a final extension. The PCR products were separated on 1.5% agarose gel and stained with ethidium bromide, followed by densitometry measurement (using image processing ImageJ program from GitHub). The amplicon of the β-actin (Hs99999903_m1) is 196 nt with the forward (F) and reverse (R) primers of F: CATGCCCACCATCAGCCCTGG and R: ACAGAGCCTCGCCTTTGCCGA, respectively. For DJ-1 (376 nt) we used F: GCCTGGTGTGGGGCTTGTAA and R: GCCAACAGAGCAGTAGGACC

### 2.4. RNA-Seq

Total RNA was extracted using the RNeasy Plus Universal Mini Kit (QIAGEN), according to the manufacturer’s protocol. Total RNA samples (1 μg RNA) were enriched for mRNAs by pull-down of poly (A). Libraries were prepared using a KAPA Stranded mRNA-Seq Kit, according to the manufacturer’s protocol, and sequenced using Illumina NextSeq 500 to generate 85 bp single-end reads (a total of 25–30 million reads per sample).

### 2.5. Bioinformatic Analysis

Next-generation sequencing data underwent quality control using FastQC, version 0.11.9. They were then preprocessed using Trimmomatic and aligned to the reference genome GRCm38 with the STAR aligner using default parameters. Genomic loci were annotated using GENCODE version M25. Genes with low expression were filtered out of the dataset by setting a threshold of a minimum of two counts per million in three samples.

Differential analyses were performed on all experimental groups, and genes with an FDR <0.05 were considered. Only genes with an absolute log fold-change of ≥0.5 were labeled up- or downregulated all the rest were considered unchanged. We partition genes by the type as coding and non-coding (including pseudogene, anti-sense, miRNAs, TEC, lncRNA and other rare biotypes).

Functional analysis and network view was based on STRING protein-protein interaction (PPI) platform. The minimal PPI confidence score ranges from 0.4 to 0.9. For protein interacting, we collected data from a human centric view from IntAct, limiting DJ-1 interacting proteins to a minimal MI score >0.6). Only to evidence of physical interactions were included [31]. We used the gene expression normalization nTPM to compare the expression of DJ-1 across cell types. Data is available in Human Proteome Atlas (HPA) [32].

### 2.6. Statistical analysis

Principal component analysis (PCA) was performed using the R-base function “prcomp”. EdgeR was used to perform RNA read counts by the trimmed mean of the M-values normalization of RNA (TMM) and differential expression analysis [33]. Figures were generated using the ggplot2 R package [34].

## Results

### DJ-1 response to oxidative stress by translocation to nucleus

PARK7 (Dj-1) was extensively studied for its function as oxidative stress sensor and in executing the oxidative stress cellular response. This role was associated mostly with neurons that are over-sensitive to oxidation insults. However, all cells and the majority of cell lines express DJ-1 at moderate to high levels (Supplementary Fig. S1). The current notion is that the modification of DJ-1 on cysteine residue, is a prerequisite for DJ-1 nuclear function, including its role in cancer and metastasis.

To this end, we tested SH-SY5Y neuroblastoma cells as a model for neuronal-like cells [35]. **Fig. 1A** shows representative frames of immunofluorescence staining with DJ-1 antibodies at different time from exposing cells to 100 μM of superoxide (H_2_O_2_). Following 24 h there is no sign for the oxidation wave of DJ-1 in SH-SY5Y cells. Maximal effect of DJ-1 entering the nucleus was achieved within 30 min (**Fig. 1B**). The kinetics of DJ-1 translocation from cytosol to nuclei dictate the downstream activation of transcription and signaling cascades mediated by DJ-1 [1].

**Figure 1.**
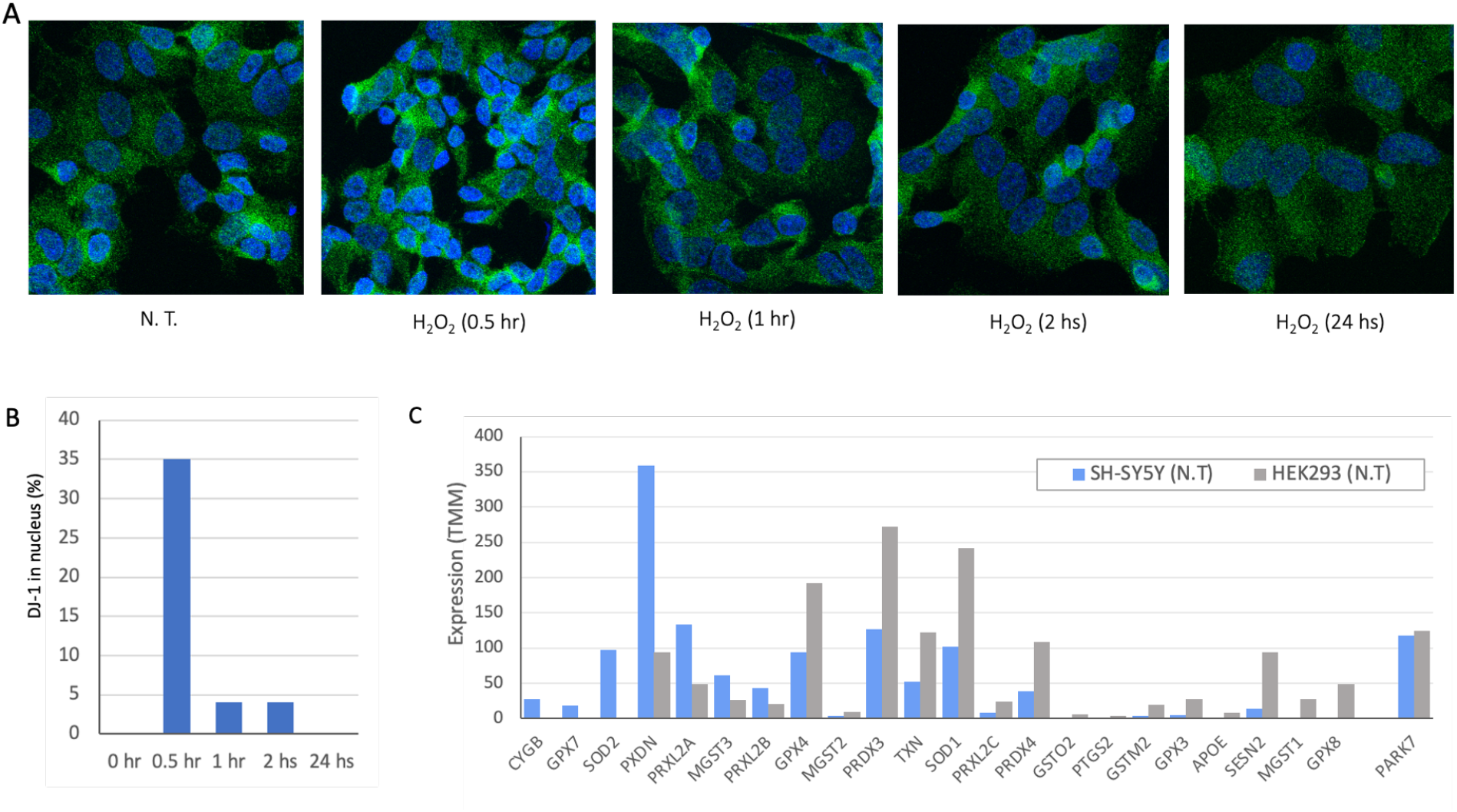
Cell lines characterization in view of DJ-1 and antioxidant activity. **(A)** Staining of SH-SY5Y cells with antibody for DJ-1 (green) and nucleus (DAPI, blue). The cells were treated with 100μM H_2_O_2_ and were fixed for immunofluorescence analysis at the time indicated from the addition of hydrogen peroxide. **(B)** Measuring the percentage of DJ-1 intensity entering the nuclei was calculated relative to non-treatment cells that were defined as 0% of DJ-1 in the nucleus. In untreated cells, we considered the cytosol and mitochondria to account for 100% of the staining. **(C)** Gene expression in cell cultures of SH-SY5Y (gray) and HEK293 (blue) cell lines for selected genes associated with antioxidant activity (76 genes, GO: 0016209). Full list is available in Supplementary **Table S1**.

We compared the genes that act in the antioxidant response in SH-SY5Y and HEK293 cells (**Fig. 1C**). There are 76 antioxidant activity annotated genes (GO: 0016209; Supplementary **Fig. S2**). The expression level of these genes in SH-SY5Y and HEK293 are shown (Supplementary **Table S1**). **Fig. 1C** shows representative genes, while the expression of DJ-1 is similar in both cell types, many of the genes (e.g., GPX8, GPX4) are highly expressed in HEK293 but display low expression levels in SH-SY5Y. Interestingly, both cell lines expressed the cytoplasmic enzyme SOD1 to different levels, only SOD2 is expressed in SH-SY5Y cells. SOD2 is located in the mitochondrial matrix and thus impacts mitochondrial apoptosis and the redox function. We anticipate that each cell displays a unique collection of genes to cope with oxidation, rendering a cell-specific capacity for coping with oxidative stress [35].

### Knockdown resulted in 8-fold suppression of the native PARK7/DJ-1 transcript

We transfected HEK293 cells by siRNA as a negative control (using esiRLUC, see Methods) and the siRNA methodology for PARK7 (see Methods). The cultured cells were collected 24 h following transfection, and total RNA (>200 nt) was prepared for RNA-seq. **Fig. 2A** shows the results of the RT-PCR at two time points (26 h and 48 h, post transfection). It is evident that already 26 h post-transfection DJ-1 transcripts are strongly suppressed. Quantitation of the degree of suppression by the specific siRNA relative to the non-specific control siRNA showed 8-fold reduction which was constant along 48 h. Note that the RULC control showed no reduction in DJ-1 transcripts, supporting the efficiency of the knockdown (KD) protocol used.

**Figure 2.**
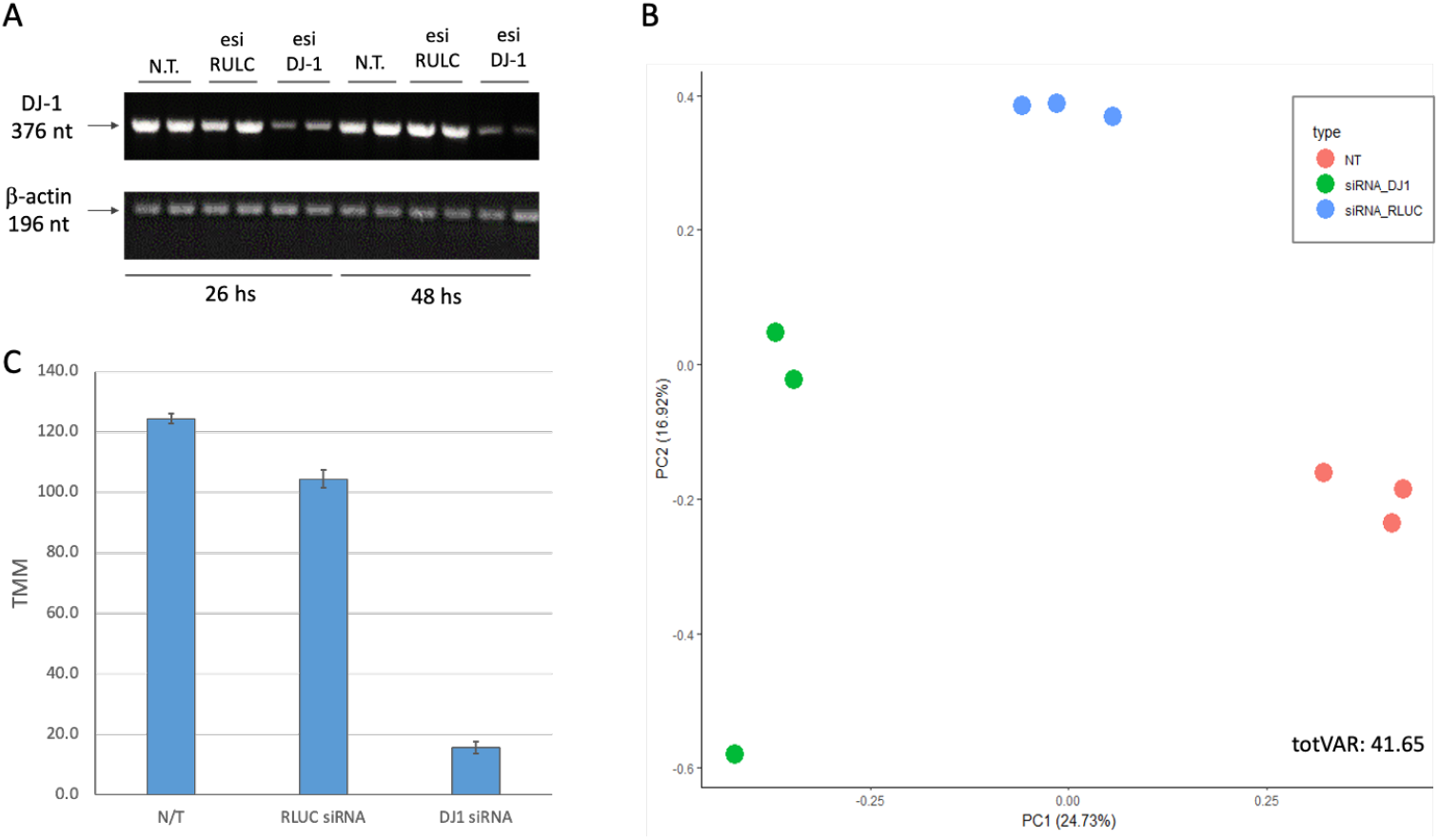
Knockdown of PARK7/DJ-1 using siRNA. **(A)** RT-PCR of the cells prior to treatment, following RULC and PARK-7 siRNA. PCR amplicons were separated by agarose gel separation. Each sample was compared to β-actin, a normalization control (196 nt). The siRNA was tested for 26 and 48 h following transfection. **(B)** PCA for 9 cell samples, based on the top 1000 differentially expressed genes (DEG) colored by the three experimental groups. The nontreated cells (N.T.), negative control transfection with esiRNA-RULC (RULC), and cells transfected with esiRNA-PARK7 (DJ-1). Each sample is represented by a colored dot. The variance explained are indicated for PC1 and PC2. The explained variance of PC1 and PC2 reached 41.6%. **(C)** The results from B for the level of expression of DJ-I is shown. Each sample was normalized by TMM methods for 1 M reads accounted for 18,158 transcripts.

**Fig. 2B** shows the results using dimensional reduction by principal component analysis (PCA). Each experimental group is represented by three biological triplicates. The PCA resulted shows the nontreated cells (N.T.), mock transfection with esiRNA-RULC (depicted RULC), and cells transfected with esiRNA-PARK7 (DJ-1). The two major components (PC1 and PC2) explained together 42% of the variance (**Fig. 2B**). The partitioned of the nine samples according to the treatments validates the quality of the results. For the PCA (**Fig. 2B**) we have included all 18k identified transcripts from RNA-seq results (Supplementary **Table S2**).

Using the results from RNA-seq with biological triplicates we calculated the extend of the knockdown of PARK7. Only 12% of the original basal levels was detected 24 h after introducing the siRNAs (**Fig. 2C**).

### Knockdown of PARK7/DJ-1 resulted in coding and non-coding differential gene expression

In analyzing all three experimental groups, we observed a large number of differentially expressed genes. The RNA-seq results reported on 18,158 transcripts, 23% of them are non-coding RNAs (total 4192). Only 6% of these transcripts met the statistical threshold of FDR <0.05, resulting of 1083 transcripts. Among these reliable results, transcripts that were differentially expressed by log2(FC) > 0.5 (i.e., >|1.412|fold) were labelled as upregulated and downregulated transcripts. The numbers of these transcripts are shown in **Fig. 3A**. The entire expression list is normalized by TMM (total 1 million).

**Figure 3.**
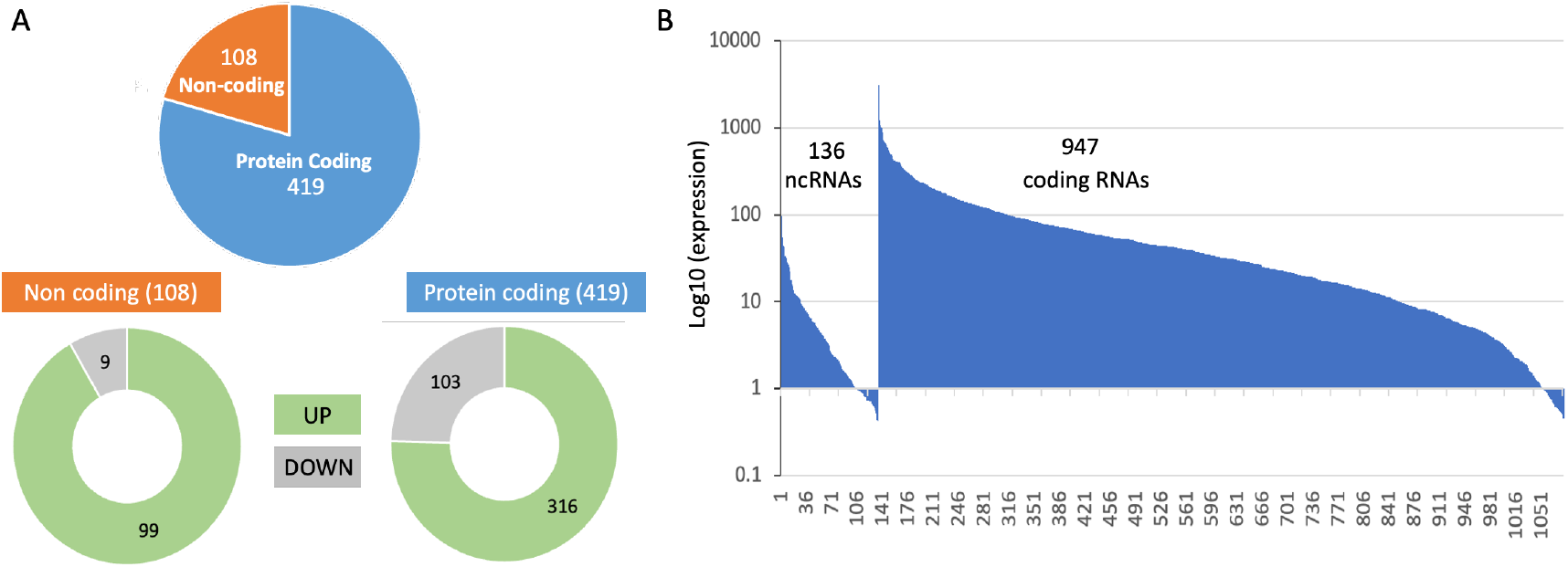
Quantitative summary of the differentially expressed transcripts of DJ-1 siRNA relative to control. **(A)** Partition of the significant differentially expressed genes (DEG) that meet the threshold of FDR <0.05 and log2(FC)|>0.5|. Partition of the DEG to those which are upregulated (UP) and downregulated (DOWN) are shown for each of the subset of protein coding (blue) and non-coding set (orange). **(B)** Ranking of 1083 DEG that are statistically significant (FDR<0.05). Genes were ranked by their expression levels with 136 ncRNAs (1-136), and additional 947 coding RNAs. The expression levels were normalized by TMM where all identified 18,158 transcripts were normalized to 1 million.

Out of the 1083 significant transcripts (**Fig. 3B**), all high expression transcripts belong to the coding genes, with 16 coding RNAs have >500 normalized TMM (e.g., actin gamma, ribosomal proteins, ADP ribosylation factor, helicase, mitochondrial membrane translocase). Among low expression gene set (209 with TMM <5.0) over a third of the them belon to non-coding RNAs, in accord with the relative lower expression levels on non-coding RNAs. The rest of the analysis will focus of coding protein genes. The RNA-seq significant results are listed in Supplementary **Table S1**.

### Non-specific siRNA induces interferon signaling and the antiviral response

Upon transfection the HEK293 with siRNA of RULC, we observed a and coordinated induction of genes that are associated with the innate immunity and more specifically with interferon response. This observation most likely reflect non-specific induction of cells to the delivery system for the duplex of RNA molecules and the activating the dsRNA pathway of viral infection. Testing the induction of esiRULC versus naïve, non-treated cells shows the upregulated set of coding transcripts (Total 75 gene, marked interferon stimulation, **Fig. 4**). The network is indicative of the activation of type I Interferons (IFN) that acts through the activation of JAK/STAT signaling cascade to trigger antiviral response. Note that STAT1 acts as a hub to several transcription factors that belong to ATF, EGR and FOS families. The ATF3 is a regulator that links inflammation, oxidative stress, and immune responses [36].

**Figure 4.**
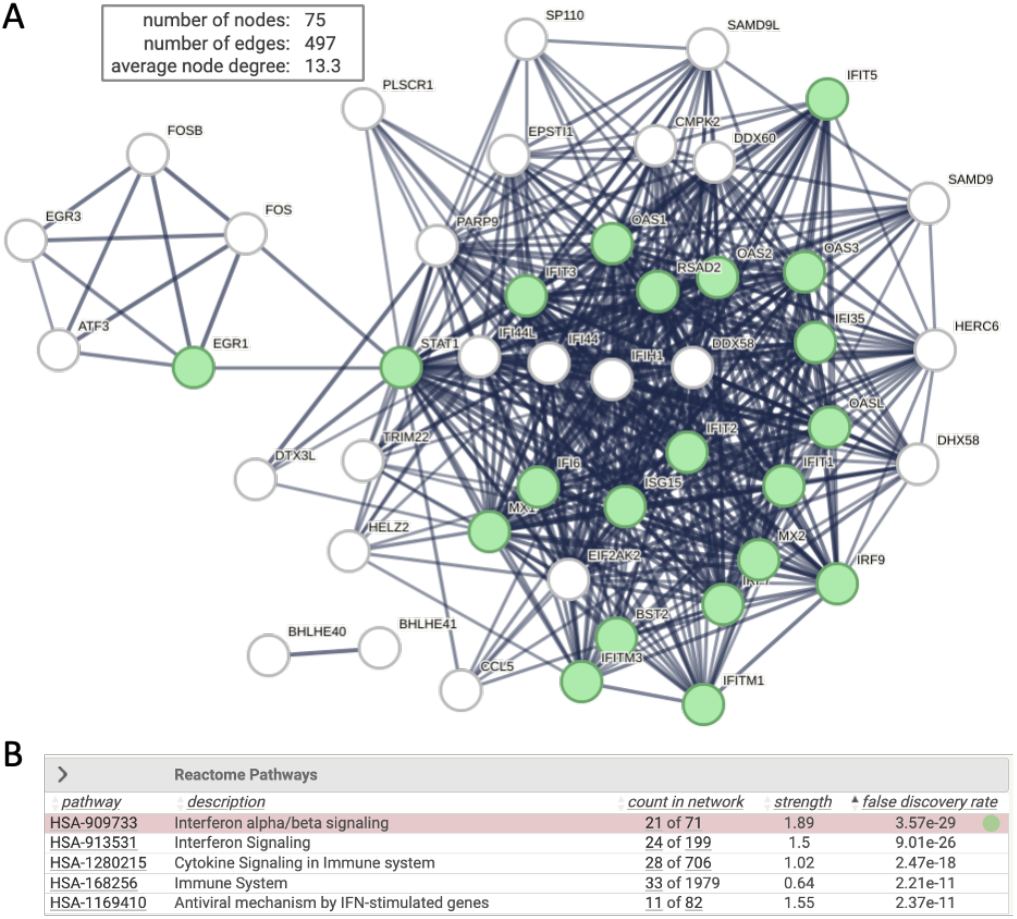
Network of interferon signaling induced by non-specific siRNA. **(A)** STRING view for 75 significantly upregulated DEG (FDR <0.05 and log2(FC) >0.5) from analysis of siRNA RULC vs. non treated (N.T.) cells. STRING PPI confidence is > 0.9. **(B)** several significant Reactome pathways are enriched. The nodes in the graph of Interferon alpha/beta signaling (HAS 909733) are colored green. The rest of the pathways are strongly connected to the immune system and antiviral IFN-induced mechanism.

### Knockdown of PARK7/DJ-1 strongly boosted the antiviral response

**Fig. 5A** shows a Venn diagram of the experiment groups. It shows results from comparing KD of DJ-1 vs. the background of siRNA of RULC and non-treated cell. Among the genes that were changed with esiRNA RULC negative control, 45 of the them also induced by using esiRNA for DJ-1.

**Figure 5.**
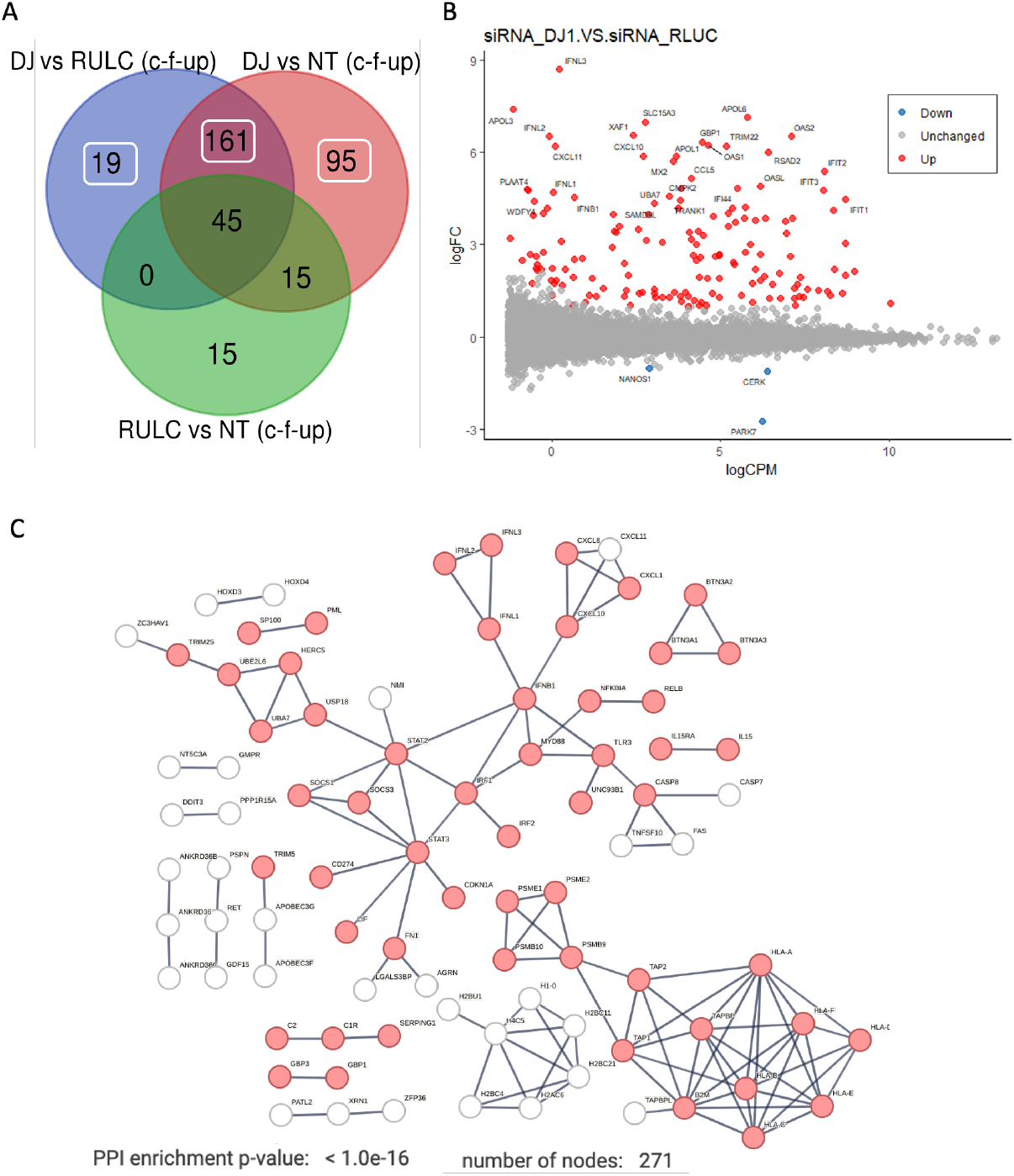
Differentially expressed transcripts of DJ-1 siRNA relative to control. **(A)** A Venn diagram of the experiment groups. The number of overlapping genes is indicated for comparing KD of DJ-1 vs. siRNA of RULC and vs. non-treated cell. The mark of c-f-UP indicates the use of filters by coding genes (c), FDR (<0.05, f) and only upregulated DEG (UP). **(B)**. MA plot is shown as the log2(FC) for each gene versus its mean expression between the two groups (y-axis) of KD of DJ-1 compared to siRNA of RULC (x-axis) Blue points indicate those that are downregulated and in red are overexpressed genes (FDR<0.05, log2(FC)|0.5|). **(C)** STRING view for 275 significantly upregulated DEG (FDR <0.05 and log2(FC) >0.5) from analysis of siRNA RULC vs. non treated (N.T.) cells. STRING PPI confidence is > 0.9. Partition of the significant differentially expressed genes (DEG) that meet the threshold of FDR <0.05 and log2(FC) of |>0.5|.

**Fig. 5B** shows an MA plot of the differential expressed transcripts. Highlighted are set genes that meet the threshold for FDR and FC. Inspecting of these genes emphasizes the occurrence of viral-like induction that span all levels of expression (y-axis). Notably, many of the genes that were strongly induced (>10 folds). Among the genes with strongest induction are the interferon (INF) genes and their modulators. For example, the set of OAS genes that recognize dsRNA as pathogen-associated molecular pattern (PAMP). The induction of OAS1, OAS2, OASL which as main sensors for viral infections explain the induction of numerous interferon induced proteins (e.g., IFIT1, IFIT2, IFI44, MX1, MX2). This coordinated signature argues for further amplification of the antiviral response following KD of DJ-1. **Fig. 5C** shows the strong connectivity of the induced genes using STRING analysis. The analysis used a filtered set by removal the non-specific set (45 genes, as in **Fig 5A**) and including the rest of the 275 genes that were upregulated as a result of KD by siRNA of DJ-1 (Supplementary **Table S3**).

The protein-protein interaction (PPI) network is composed of several components: (i) the antigen presenting that include genes such as TAP1 and TAP2 that together form the TAP protein complex (transporter associated with antigen processing). (ii) the histone with H2 and H4 genes that are indicative of epigenetic, gene expression and chromatin reorganization. (iii) the main component of interferon and the innate immune response. The hub proteins include the STAT3, IRF1 and INFB1 genes. Specifically, the IRF1 (interferon regulatory factor 1) is a transcription factor and activator of interferons alpha and beta and gamma. In addition, the IRF1 plays a role in apoptosis and tumor-suppression. The result of the activation of type I interferon is shown in the network (**Fig. 5C**) by the increase in the inflammatory cytokines and chemokines. Note the activation of smaller connected components such as the BTN (butyrophilin) gene set which belong to the major histocompatibility complex (MHC)-associated genes, and genes from the classical pathway of the complement system (e.g., C1, C2).

### Knockdown of PARK7/DJ-1 exposed downregulation of genes with membrane-associated functions

We identified a small set of genes that were substantially downregulated along with the KD of DJ-1 (57 genes). There is no sign for involvement of the innate system among the downregulated genes (**Fig. 6A**). We noted that the degree of downregulation in modest. Inspecting the network shows that on average, the node degree is low, even at a relaxed threshold of PPI confidence (score >0.4). The gene that was maximally downregulated is CERK (ceramide kinase; 2.2 folds; Supplementary **Table S3**). CERK is associated with ceramide metabolism through the effect of ceramide and sphingolipids on mitophagy [37]. Another downregulated gene is GET1 that determines the kinetics of mitochondrial targeting proteins, therefore also control mitophagy [38]. Surprisingly, among the downregulated genes (Supplementary **Table S2**) many were assigned to neurons (NDNF, neuron derived neurotrophic factor; NANOS1, nanos C2HC-type zinc finger 1) and synapses, including synaptic vesicle proteins (e.g., synaptophysin, SYP). We attribute this observation to the role of these genes in membrane trafficking. For example, syntaxin 6 (SNX6) functions as an endosomal organizer and intracellular protein transport. Another neuronal-like gene is SYT16, a calcium-independent membranous protein is involved in trafficking of secretory vesicles in non-neuronal tissues. Other genes were implicated in controlling neuronal survival and differentiation (e.g., GFRA1, a member of the glial cell line-derived neurotrophic factor (GDNF) receptor family), and cellular adhesion (CNTNAP2 that mediates interactions between neurons and glia during during development). Other genes belong to the TNF-receptor superfamily (e.g., TNFRS10D, TNFRSF21) indicative for their role in inflammation and in inhibition of cell apoptosis.

**Figure 6.**
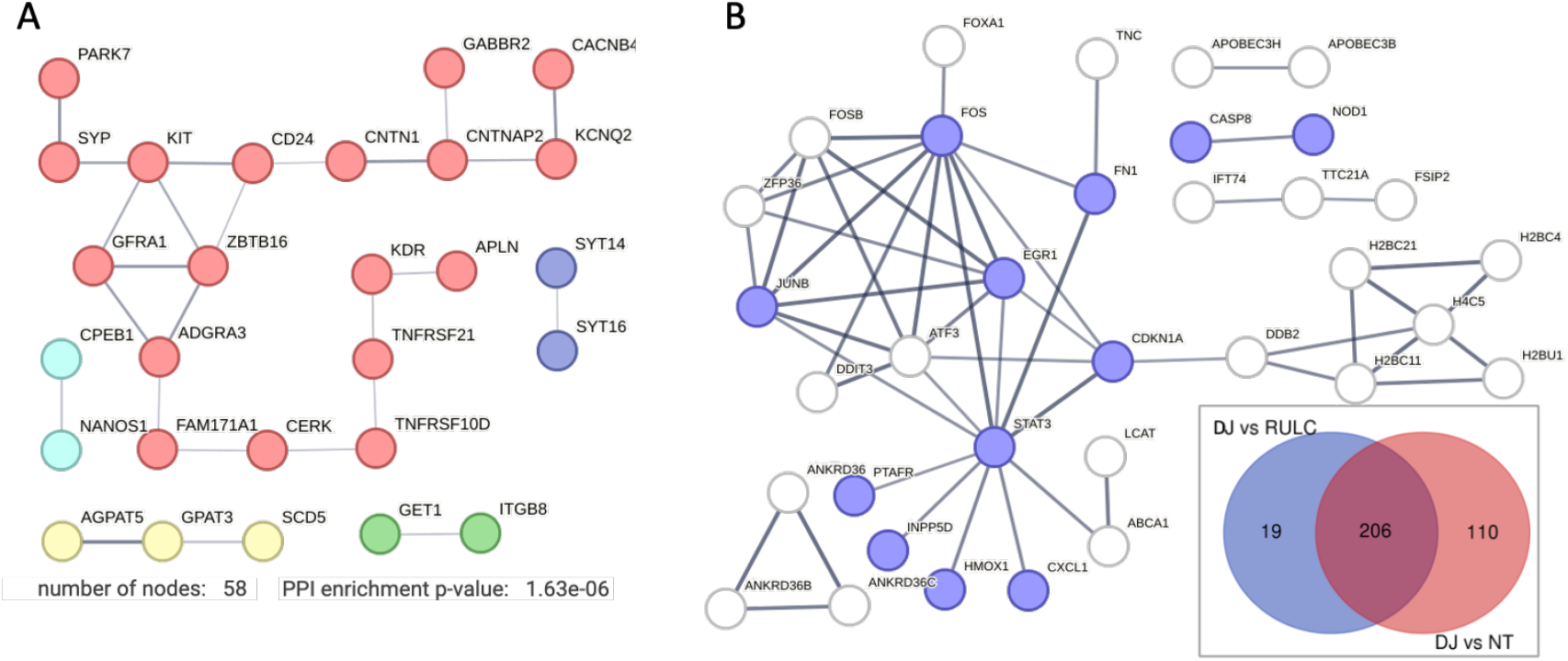
DJ-1 dependent effects on DEG. **(A)** STRING based network for all genes that were significantly downregulated with threshold of FDR <0.05 and log2(FC)<-0.5. STRING PPI confidence score >0.4. **(B)** STRING based network for 129 genes that were significantly upregulated with threshold of FDR <0.05 and log2(FC)>0.5 (inset, Venn diagram of KD of DJ-1 compared to siRNA of RULC and N.T. cells; 19 and 110 genes). STRING PPI confidence score >0.7. The genes in purples are enriched genes annotated by Reactome pathway HSA-1280215: cytokine signaling in immune system (q-value 0.0097).

**Fig. 6B** indicates the set of genes that can be attributed to the KD of DJ-1 after filtering out the 206 upregulated overlapping genes (inset, Venn diagram). We focused on 129 upregulated DJ-1 gene set (**Fig. 6B**). These genes defined as significant (PDR <0.05) among coding genes that showed a minimal log (FC) >0.5). We identified STAT3 as a hub that connect stress, inflammation with transcription regulation (e.g., EGR1 and ATF3) and the DNA repair and cell cycle axis (CDKNA1 and DDB2). By eliminating the overwhelming signature of antiviral response, we could highlight smaller gene set that were induced. As example is the set of ANKRD36 family members, there were implicated in the pathology of immune and metabolic diseases [39], and considered as potential diagnosis marker of Chronic myeloid leukemia (CML) [40].

## Discussion

In this study, we investigated the role of KD in DJ-1 in the cellular context of HEK293 cells. We identified a robust and strong antiviral response that was moderately induced by using the non-specific siRNA of RULC. The use of siRNA methodologies across different cell lines is a variable that is poorly controlled. Most commercially used siRNA protocols rely on a combination of a small set of specific sequences (about 4), while the negative control setting uses scrambled sequences to ensure specificity. Several systematic studies indicated the importance of the cell type-dependent nature of sensitivity to siRNA by its length and its impact on gene expression related to the innate immune response. For example, in HeLa cells, introducing dsRNAs longer than 23 bp induced toxicity and cell death, while shorter RNA fragments did not affect viability. In all cell types, the short siRNA duplex (19 nt) did not affect cell viability [41].

It is known that siRNA delivery reagents may induce stress in the subjected cells. Specifically, Lipofectamine 2000 (Lipo2000) is characterized by high transfection efficiency. As a byproduct of the use of Lipo2000 for siRNAs, autophagy is induced to cope with cellular oxidative and ER stress signals [42]. We showed a strong induction of the antiviral response in HEK293 with no sign of cytotoxicity, and we still achieved effective silencing with only 12% of the basal level of expression remaining following 24-48 h post transfection of siRNAs. The siRNA system and delivery are also sensitive to the presence of sequence motifs via activation of toll-like receptors (e.g., TLR8) to induce an undesirable antiviral response [43]. Due to the importance of siRNA for research and as a promising therapeutic method, optimization is needed to achieve the desired responses while optimizing silencing efficiency.

Similar to our finding, siRNAs (21 nt) that were introduced into cells for silencing specific genes, triggered the JAK/STAT signaling pathway [44]. The signaling cascade was initiated by EIF2AK2 (PKR, an IFN-induced dsRNA-dependent kinase), which plays a central role in the innate immune response to viral infection [45]. In our experiments, EIF2AK2 was already overexpressed (4 folds) by the siRNA of the RULC, which is likely to mediate the observed antiviral response.

It was shown that DJ-1 deficiency or knockdown in primary cortical neurons and mouse embryonic fibroblasts caused alternations in mitochondrial shape [46]. Knockout mice and DJ-1 pathogenic mutations led to elevated ROS production, lower mitochondrial membrane potential, and changes in gene expression, which can thus explain the importance of DJ-1 function in neurodegenerative diseases [47]. These in vivo findings corroborate the observed effects of suppressing DJ-1 on trafficking, mitophagy, and redox homeostasis (**Fig. 6A**).

We observed that the antiviral effect was amplified by KD of DJ-1 (with ∼200 overexpressed genes and ∼100 non-coding RNAs). While no function is known for most of these non-coding RNAs, the upregulation of lncRNA of NEAT1 (2.2 folds, q-value 3.7e-05) is intriguing, as NEAT1 was proposed to be upregulated in response to oxidative stress [48] and as a potential biomarker in Parkinson’s diseases [49]. We questioned whether the effect of KD of DJ-1 is through signaling convergence of the IFN signaling pathways or, alternatively, through the failure of the cells to cope with the alteration in the change in redox level. We suggest that it is the former explanation. We have not challenged the cells by an oxidative stimulus [35], and as anticipated, no sign of cytotoxicity following siRNAs was observed, which argues for uninterrupted mitochondrial homeostasis. In contrast, the functional networks presented in this study (**Figs. 4**–**6**) indicated the key role of JAK/STAT as the main hub. DJ-1 negatively regulates inflammatory responses in astrocytes and microglia by facilitating interactions between STAT1 and its phosphatase, SHP-1 [23]. Experiments performed on DJ-1 knockout mice led to increased inflammatory mediators compared to wild-type mice. This was explained by the direct interaction of DJ-1 with SHP-1 and shifting the levels of phosphorylated STAT1. Therefore, DJ-1 may prevent excessive STAT1 activation and reduce the risk of brain inflammation [23].

In this study, we tested the KD of DJ-1 in non-neuronal cell lines by inspecting the transcriptional response. We emphasize that different cells present different sensitivity to various stress and external stimuli (**Fig. 1**). Different cells substantially differ in their capacity to respond to molecular perturbations and external stimuli [50]. We argue that the depletion of DJ-1 in HEK293 cells caused a slight alteration in membranous trafficking (**Fig. 6B**) but mostly a change in transcription, rendering the cells labile and prone to multiple stressors, including dsRNA-INF dependent stress. Our study highlights the intricate interplay between siRNA technology and the robust activation of the innate

## Author Contributions

Conceptualization M.L and K.Z..; Methodology, K.Z. and M.L.; Software, K.Z.; Formal analysis, K.Z. and M.L.; Resources, K.Z.; Writing the original draft, M.L. Visualization, K.Z. and M.L.; Project administration, M.L.

## Informed Consent Statement

All authors share their consent for publication.

## Data Availability Statement

RNA-seq data files are available through the supplemental tables and Figures.

## Acknowledgments

We would like to thank the members of Linial’s lab for useful comments throughout the research project. Special thanks to Dr. Tsiona Eliyahu and Eliran Giladi for their help in the early phase of this research and to Prof. Michal Goldberg lab for fruitful discussion.

## Abbreviations

AD: Alzheimer’s disease
BSA: Bovine serum albumin
CNS: Central nervous system
DEG: Differentially expressed genes
DMEM: Dulbecco’s modified Eagle medium
FC: Fold change
FDR: False discovery rate
HPA: Human proteome atlas
GO: Gene ontology
IFN: Interferon
NT: Not treated
PCA: Principal component analysis
PCR: Polymerase chain reaction
PD: Parkinson’s disease
RNA-seq: RNA sequencing analysis
RT: Reverse transcription
TMM: Trimmed mean of means

