## Supplemental Fig. S1-S2 for "Knockdown of DJ-1 Resulted in a Coordinated Activation of the Innate Immune Antiviral Response in HEK293 Cell Line"

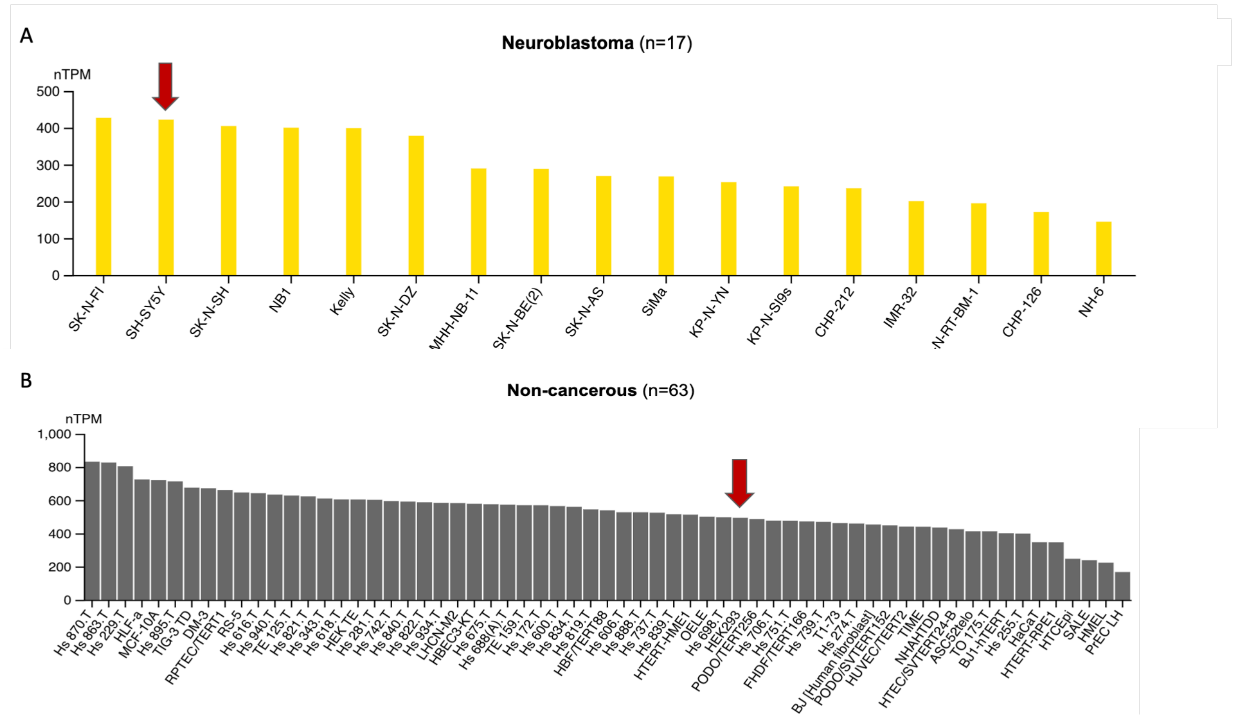


**Figure S1.** PARK7 expression levels extracted from neuroblastoma and cancerous cell lines. Data was extracted from HPA (see Methods). Experimental data was normalized and harmonized to allow confident comparing different cells within and across groups (using nTPM). **(A)** The expression levels of the SH-SY5Y among neuroblastoma cell line (17 cells, 423 nTPM). (B) PARK7 expression levels in HEK293 is 493.5 nTMP. The range of expression among the 63 non-cancerous cell lines is 4-5 folds. Red arrows mark the cells of interest.

Figure S2.

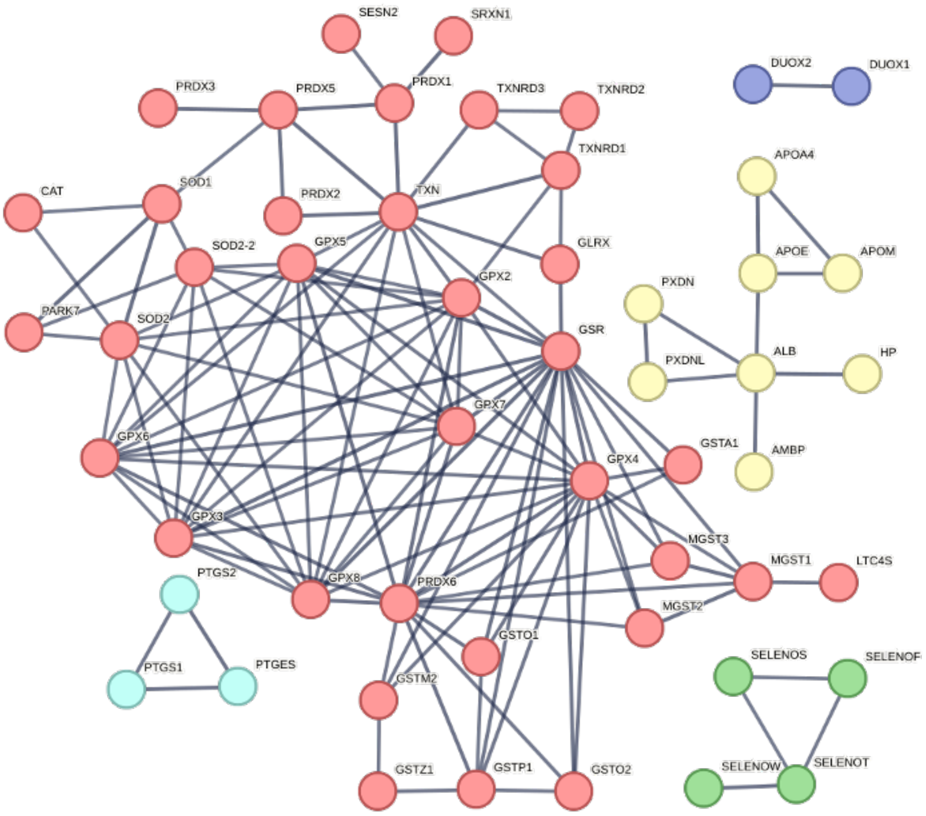


| bubble | cluster Id | gene count | protein names |
|---|---|---|---|
| | Cluster 1 | 35 | CAT, GLRX, GPX2, GPX3, GPX4, GPX5, GPX6, GPX7, GPX8, GSR, GSTA1, GSTM2, GSTO1, GSTO2, GSTP1, GSTZ1, LTC4S, MGST1, MGST2, MGST3, PARK7, PRDX1, PRDX2, PRDX3, PRDX5, PRDX6, SESN2, SOD1, SOD2, SOD2-2, SRXN1, TXN, TXNRD1, TXNRD2, TXNRD3 |
| | Cluster 2 | 8 | ALB, AMBP, APOA4, APOE, APOM, HP, PXDN, PXDNL |
| | Cluster 3 | 4 | SELENOF, SELENOS, SELENOT, SELENOW |
| | Cluster 4 | 3 | PTGES, PTGS1, PTGS2 |
| | Cluster 5 | 2 | DUOX1, DUOX2 |

**Figure S2.** Gene set of antioxidant activity (76 genes, GO: 0016209). A STRING-based network of the connected set of “antioxidant activity” (STRING PPI score >0.9). The genes are partitioned into 5 colored groups (Top), with a dominant cluster (red; 35 genes). A full list is all listed genes is in Supplementary **Table S1**.
